# Development of Surface Enhanced Raman Spectra coupled with Machine Learning Analysis for Differentiation of Closely Related Species within *Enterobacter cloacae* Complex

**DOI:** 10.1101/2025.05.08.652998

**Authors:** Menglan Zhou, Xuesong Xiong, Yanbing Li, Yingchun Xu, Bing Gu

## Abstract

**Background:** *Enterobacter cloacae* complex (ECC) is an important nosocomial pathogen and consists of multiple similar species. The taxonomy of ECC has been consecutively updated, adding to its identification difficulty.

**Methods:** A total of 92 ECC strains isolated from bloodstream infections during 2015-2020 were collected from a tertiary hospital in China. All the strains were identified by Vitek 2 Compact and Vitek MS and then subjected to whole genome sequencing (WGS) for average nucleotide identity (ANI) analysis. Surfaced-enhanced Raman spectroscopy (SERS) with a set of machine learning algorithms was applied in identifying species within ECC.

**Results:** Seven species were identified through ANI, including 28 *E. hormaechei* subsp. *steigerwaltii*, 17 *E. hormaechei* subsp. *xiangfangensis*, 12 *E. cloacae*, 11 each of *E. hormaechei* subsp. *hoffmannii* and *E. bugandensis*, seven *E. kobei* and six *E. roggenkampii*. The Vitek 2 compact indistinguishably identified all the strains as ECC and Vitek MS correctly identified one strain of *E. kobei* while achieving ambiguous results for all the other isolates. SERS combined with XGBoost model achieved 97.75% accuracy with an area under the ROC curve value of 0.9982 in the identification of ECC.

**Conclusion:** SERS coupled with machine learning algorithms holds a promising potential to acquire early prediction of ECC, outperforming the capabilities of other methods.

## 1. Introduction

*Enterobacter* spp. is one of the notorious “ESKAPE” bugs (*Enterococcus faecium, Staphylococcus aureus, Klebsiella pneumoniae, Acinetobacter baumannii, Pseudomonas aeruginosa*, and *Enterobacter* species), among which the the *E. cloacae* complex (ECC) is of major importance (Miller & Arias, 2024). *Enterobacter* spp. consists of closely related species that cannot typically be identified precisely by common phenotypic tests (Mezzatesta et al., 2012). Moreover, the taxonomy of genus is complicated by the reassignment of some species to other genus. For example, *E. aerogenes* has been moved to genus *Klebsiella* (Tindall et al., 2017), *E. agglomerans* to genus *Pantoea* (Gavini et al., 1989), and *E. sakazakii* to genus *Cronobacter* (Iversen et al., 2008). ECC represents the most frequently isolated *Enterobacter* spp. in human respiratory, urinary tract, and bloodstream infections, especially in immunocompromised individuals (Mezzatesta et al., 2012; Wisplinghoff et al., 2004). Based on available epidemiology data, ECC has become the third major drug-resistant Enterobacteriaceae species involved in nosocomial infections after *Escherichia coli* and *Klebsiella pneumoniae* (Zhou et al., 2018). According to China Antimicrobial Surveillance Network (CHINET) data, ECC resistance to carbapenems increased from 4.8% in 2010 to 9.7% in 2022, and the NDM carbapenemase is mostly detected in China which can not be inhibited by current enzyme inhibitors (Giamarellou, 2016; Zong et al., 2021). The resistance to polymyxins, another resort to CRE, was also increasing, from 2% to 5% from 2019 to 2022, even higher than *Escherichia* and *Klebsiella* (usually <2% for both) (Giamarellou, 2016; Zong et al., 2021). What’s more, ECC exhibited high and variable heteroresistance to polymyxins (Fukuzawa et al., 2023; Guerin et al., 2016).

Given the diverse nature of species and resistance distribution among different ECC, it is of great significance to precisely identify ECC into species and subspecies levels. Phenotype-based identification methods such as commercial automated biochemical assays and matrix-assisted laser desorption/ionization time-of-flight mass spectrometry (MALDI-TOF MS), have been commonly used in clinical microbiology laboratories, but often failing to differentiate the species within ECC. The application of whole-genome sequencing (WGS) enables the precise identification of ECC through average nucleotide identity (ANI) and DNA-DNA hybridization (DDH). However, both methods rely on WGS which is relatively high-cost and inconvenient for routine clinical labs. Surface enhanced Raman spectroscopy (SERS) is an emerging technique based on interactions between the light and chemical bonds (Tang, Li, et al., 2022). In recent years, the potential application of SERS in bacterial pathogen detection has been extensively explored, especially in certain closely related species, such as the *Shigella* spp. and *Escherichia coli*, the *Acinetobacter baumannii*/*calcoaceticus* complex (Liu et al., 2023; Xiong et al., 2024).

Being one of the clinical important pathogens, so far, the usage of SERS for the identification of ECC has never been investigated. In this study, we firstly applied the SERS technique combined with machine learning models for rapid and accurate discrimination of ECC.

## 2. Methods and materials

### 2.1 Strains collection and identification

A total of 92 consecutive ECC strains isolated from bloodstream infections during 2015-2020 in Peking Union Medical College Hospital (PUMCH) were included in this study. All the strains were initially identified by Vitek 2 Compact and Vitek MS (bioMérieux, France) and then subjected to WGS using a paired-end library with an average insert size of 350 bp (ranging from 150 to 600 bp) on a HiSeq sequencer (Illumina, USA). Sequence quality was evaluated using fastqc and fastp, the N-base, Q-value<30 and poor-quality reads were removed. Sequence was assembled with SPAdes (SPAdes-3.15.5). The quality after assembly was evaluated and processed with Trimmotic and seqkit; contigs shorter than 200 bps were removed from the results. The ANI was analyzed with fastANI (Konstantinidis & Tiedje, 2005), and the type strains of *Enterobacter* species and subspecies were collected from NCBI Refseq Database (Tang, Li, et al., 2022). The classification criteria adhere to ANI > 95% for species and ANI > 98% for subspecies.

### 2.2 Synthesis of silver nanoparticle (AgNPs)

AgNPs synthesis followed the routine procedures that were previously reported by Tang et al. with modifications (Tang et al., 2023). Briefly, 33.72 mg of Silver Nitrate AgNO_3_ (Sinopharm, Beijing, China) was dissolved into 200 mL ultra-pure water (deionized distilled water, ddH_2_O) with heating via a benchtop magnetic stirrer (ZNCL-BS230, Shi-Ji-Hua-Ke Pty. Ltd., Beijing, China) until boiling. After that, stop heating, add 8 mL of sodium citrate (1% wt) to the solution with stirring speed at 650 RPM until the mixture was cooled down to room temperature. Final volume of the solution was adjusted to 200 mL via double-distilled water (ddH_2_O). One milliliter of the solution was transferred to a 1.5 mL Eppendorf tube, which was then centrifuged at 7,000 RPM for 7 min (Centrifuge 5430 R, Eppendorf, USA). After centrifugation, the supernatant was discarded and the pellet was resuspended with 100 μL of ddH2O, which was considered as AgNPs substrate and was stored in the dark at room temperature for later use.

### 2.3 Measurement of SERS spectra

All strains of ECC were cultured overnight on Columbia blood agar plates. Then, ddH_2_O and DensiCHEK Plus (bioMérieux, France) were used to adjust the concentration of bacterial solutions to a 2.0 McFarland (6.0 ×10^8^ cells/mL). The same volume AgNO_3_ and bacterial solutions of 2.0 McFarland with 5 microliters were vigorously mixed on a vortex mixer for 5 s, 5.0 μL of which was then dropped onto a silicon wafer to dry naturally, and then 50 spots were randomly selected for automatic acquisition of SERS spectra. In particular, map image acquisition (matrix mode) of a Renishau Raman instrument was used to scan points (step = 10 mm, x = 5, y = 10). A total of 50 points were auto-scanned, and a large range was randomly selected for matrix scanning for each sample. Three independent replicates were performed for each isolate.

### 2.4 Average SERS spectra and characteristic peaks

During SERS spectral analysis, average SERS spectrum for each study group was generated by calculating the average Raman intensity at each Raman shift in the range of 500 to 1800 cm^-1^. All the averaged Raman spectra were pre-processed using LabSpec 6 (HORIBA Scientific, Japan), including smoothing, denoising, baseline correction and normalization. Characteristic peaks of each average SERS spectrum were then identified. Specific processes were performed as follows: 1) *Smoothing* function was first used to smooth and denoise the spectrum with settings of *Degree*=4, *Size*=5, and *Height*=50; 2) *Baseline Correction* was conducted through the following parameters: *Type*=Polynom, *Degree*=6, *Attach*=No, while *Baseline Fitting* was done via *Auto* function; 3) *GaussLoren* function was used to find characteristic peaks, and the parameters were set to *Level*=13% and *Size*=19 while other parameters were set to default; 4) *Normalization* function was used in default settings to normalize all the SERS spectra in order to compare the average SERS spectral curves for different sample groups; 5) *Search* function was finally used to identify characteristic peaks. Characteristic peaks for each average SERS spectrum were labelled in vertical black arrows at corresponding Raman shifts. The software Origin (OriginLab, USA) was used to generate the 20% standard error band for each average SERS spectrum. The width of the standard error band reflected the reproducibility of each SERS spectrum.

### 2.5 Machine learning analysis of SERS spectra

Due to the complexity of SERS spectral data, classical statistical methods are insufficient to analyze Raman spectral data. Therefore, machine learning algorithms have been recruited to analyze and predict the identification of SERS spectrum. To obtain an effective identification model for different EC<u>C strains</u>, we compared the performance of six ensemble learning algorithms: Adaptive Boosting (AdaBoost), Decision Tree (DT), Random Forest (RF), Support Vector Machine (SVM), Gradient Boosting (GBoost) and eXtreme Gradient Boosting (XGBoost). Before conducting machine learning (ML) analysis, we employed the *train_test_split* function to split all SERS spectral data into training, validation, and test sets in a 6:2:2 ratio. We used the *LabelEncoder* function and the *to_categorical* method in the Scikit-Learn package (version 0.21.3) to convert the sample labels in the dataset into label-encoded form. The test dataset was exclusively used to assess the predictive performance of the models and was not utilized for training and validation. During the training of the six machine learning models, we employed grid search functions to train and fine-tune model parameters. Specifically, *GridSearchCV* function was used to optimize the hyperparameter combination, and the *cv* parameter was set to 5, which means that five times of cross validation would be performed. The hyperparameter combination with the highest average score was taken as the best for the final model training.

### 2.6 Evaluation of deep learning model

To validate the ability of machine learning models to distinguish ECC, quantitative metrics were employed to assess the performance of these algorithms, such as accuracy_score (ACC), precision_score (Pre), recall_score (Recall), and f1_score (F1).The most frequently used evaluation metric is ACC, which represents the proportion of correctly predicted among all samples. ACC was calculated using the accuracy_score function. In addition, we used the “Precision” function to calculate the probability that the sample predicted by the model to be a certain strain was actually this type of strain, and the “Recall” function was used to calculate the probability of all strains of a certain type correctly identified by the model. Since “Precision” and “Recall” were a pair of contradictory quantities, the F1-Score, as the harmonic average of these two indicators, was calculated for a comprehensive evaluation. The average parameter for F1-Score is weighted. To prevent model overfitting during training, we used the 5-Fold Cross-Validation (CV) method. This involved setting cv = 5 in the *cross_val_score* function to split the training dataset into five equal-sized sub-datasets.

Except for the quantitative evaluation metrics, we also used the receiver operating characteristic (ROC) curve to visualize the performance of the model on the test set. The roc_curve and *roc_auc_score* methods in the metrics function were used to calculate the area under the curve (AUC) values. In order to further show the detailed prediction of the model on the test set data, we used the *confusion_matrix* function to calculate the probability of the predicted value and the true value of the model, and the output probability values were input into a prewritten *plot_confusion_matrix* function and visualized as a confusion matrix. The probability that the model correctly predicted each strain was calculated and displayed in the diagonal of the confusion matrix.

## 3. Results

### 3.1 Strains identification

Using a 95% and 98% ANI cutoff to define species or subspecies boundaries, all 92 ECC strains were strictly identified. *E. hormaechei* was the most common species detected (56/92, 60.9%), which can be subdivided into *E. hormaechei* subsp. *steigerwaltii* (28/92, 30.4%), *E. hormaechei* subsp. *xiangfangensis* (17/92, 18.5%) and *E. hormaechei* subsp. *hoffmannii* (11/92, 12.0%). The remaining four species were *E. cloacae* (12/92, 12.0%), *E. bugandensis* (11/92, 12.0%), *E. kobei* (7/92, 7.6%) and *E. roggenkampii* (6/92, 6.5%). On the contary, the Vitek 2 compact indistinguishably identified all the strains as ECC, with a confidence value ranging from 95%-99%. The Vitek MS correctly identified one strain of *E. kobei* with a confidence value of 99.9% while achieving ambiguous results for all the other isolates, the majority of which was misidentified as *E. cloacae* and *E. asburiae*, each with a 50.0% confidence value (Supplementary Table S1).

### 3.2 Average, deconvoluted and characteristic peaks of SERS spectra

The full Raman spectra of bacteria contain morphological characteristics and physiological information of bacteria. In this study, the SERS spectra of ECC were collected separately. We computed the average Raman spectra and standard deviations of seven ECC species, thereby quantitatively revealing the overall trends and data variations among the SERS spectra of each bacterial type. As shown in Figure 1.A-G, the SERS spectrum repeatability of the seven ECC species was good, and the reproducibility of the SERS spectra varying within an acceptable range. However, due to the morphological and physiological similarities among the bacterial cells of the seven ECC species, we also employed deconvolution techniques to generate SERS component bands directly associated with molecular structures. As can be seen from Figure 1.H-N, the deconvoluted spectra are composed of a series of *Voigt* sub-bands, where each sub-band represents a spectral characteristic peak. By deconstructing different sub-bands, the differences among the seven ECC species are amplified. This method effectively extracts the important characteristic peaks from the Raman spectra and eliminates interference from spurious peaks.

**Figure 1.**
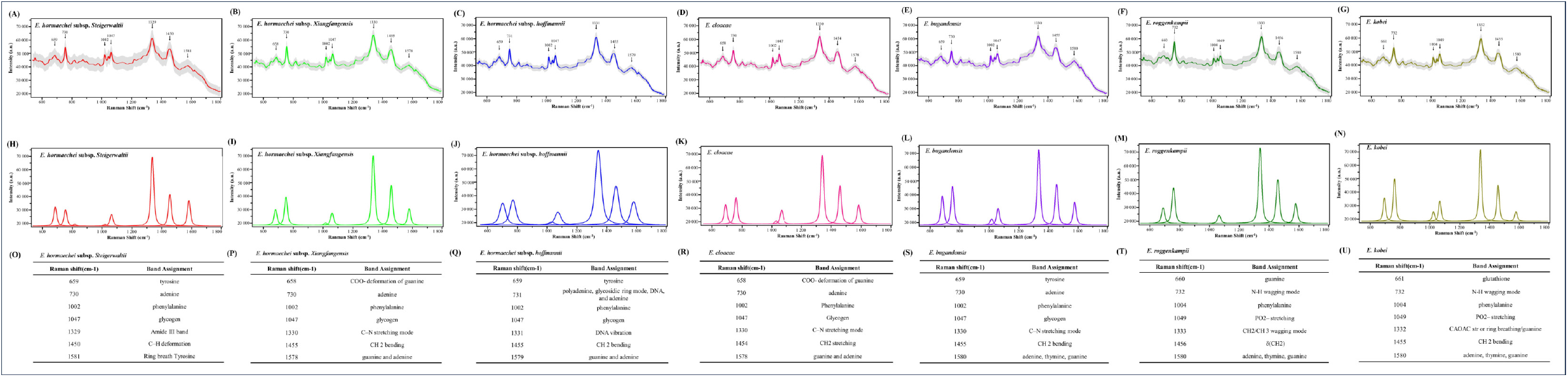
Average, deconvoluted and characteristic peaks of SERS spectra of *E. hormaechei* subsp. *steigerwaltii, E. hormaechei* subsp. *xiangfangensis, E. hormaechei subsp. hoffmannii, E*.*cloacae, E. bugandensis, E. roggenkampii* and *E. kobei*. (A-G) Average SERS spectra of ECC species. (H-N) Deconvoluted SERS spectra of ECC. X-axis is Raman shift in the range of 500-1800 cm^−1^; Y-axis is the relative Raman intensity. a.u.: artificial unit; the grey dashed area stands for the 20% standard error band for each average SERS spectrum. (O-U) Characteristic peaks of ECC species.

We also examined the characteristic peaks of different ECC species based on SERS spectra (Figure 1.O-U). Due to the high similarities within the complex, there were still some identical peaks but with significantly varied intensities. For example, the molecular vibration at 658 cm^-1^ represents COO-deformation of guanine (Ciloglu et al., 2021), appearing in both *E. hormaechei subsp. xiangfangensis* and *E*.*cloacae*, 730 cm^-1^ represents adenine (Chen et al., 2019), appearing in *E. hormaechei subsp. steigerwaltii, E. hormaechei subsp. xiangfangensis, E*.*cloacae*, and *E. bugandensis*. As for the unique characteristic peaks, they represented different molecular components and vibrations. For instance, in *E. hormaechei* subsp. *Steigerwaltii*, the peak at 1,329 cm^-1^ was associated with Amide III band (Wang et al., 2023), 1,450 cm^-1^ was associated with C–H deformation (Fan et al., 2011), and 1,581 cm^-1^ was assigned to the Ring breath Tyrosine (Yaacob et al., 2021). A prominent peak at 1,331 cm^-1^ of *E. hormaechei* subsp. *hoffmannii* was associated with DNA vibration (Walter et al., 2011). The vibration observed at 1,454 cm^-1^ in *E*.*cloacae* was attributed to CH2 stretching (Lu et al., 2011). In *E. kobei*, the unique peak at 661 cm^-1^ and 1,332 cm-1 was assigned to glutathione and CAOAC str or ring breathing/guanine respectively (Du et al., 2022; Torre-Gutierrez et al., 2022).

Based on the results above, *E. hormaechei subsp. steigerwaltii* can be uniquely identified based on characteristic peaks at 1329 cm^-1^, 1450 cm^-1^ or 1581 cm^-1^; *E. hormaechei subsp. hoffmannii* be identified based on characteristic peaks at 731 cm^-1^, 1331 cm^-1^ or 1579 cm^-1^; *E*.*cloacae* be identified based on characteristic peaks at 1454 cm^-1^; *E. roggenkampii* be identified based on characteristic peaks at 660 cm^-1^, 1004 cm^-1^, 1049 cm^-1^, 1333 cm^-1^ or 1456 cm^-1^; *E. kobei* be identified based on characteristic peaks at 661 cm^-1^, 1332 cm^-1^. *E. hormaechei subsp. xiangfangensis* can be identified based on the characteristic peaks at 658 cm^-1^ combined with 1330 cm^-1^ or 1455 cm^-1^. *E. bugandensis* can be identified based on the combination of characteristic peaks at 659 cm^-1^ plus 1589 cm^-1^.

### 3.3 Clustering analysis using Orthogonal Partial Least Squares Discriminant Analysis (OPLS-DA)

Clustering analysis was used to determine whether the seven different ECC species are separable based on their clustering in the feature coordinate system. We used the clustering method named OPLS-DA, to analyze the spectra of ECC. The results without normalization revealed that the OPLS-DA algorithm exhibit a relative lower degree of fitting between the features and spectral samples of input matrix (R2X=0.997, R2Y=0.666) and predictive capability for unknown samples (Q2=0.165) (Figure 2A). In contrast, OPLS-DA with normalization (Figure 2B) can better preserve the association rules between different spectra due to prior learning. This enables the model to exhibit a relative higher degree of fitting between the features and spectral samples of input matrix (R2X=0.980, R2Y=1.000), indicating that normalization can effectively distinguish the spectra of different ECC species. Furthmore, it also demonstrates predictive capability for unknown samples (Q2=0.313). However, overlapping of spectral data points still exists. Therefore, we need to seek more advanced machine learning methods to build rapid identification models of ECC.

**Figure 2.**
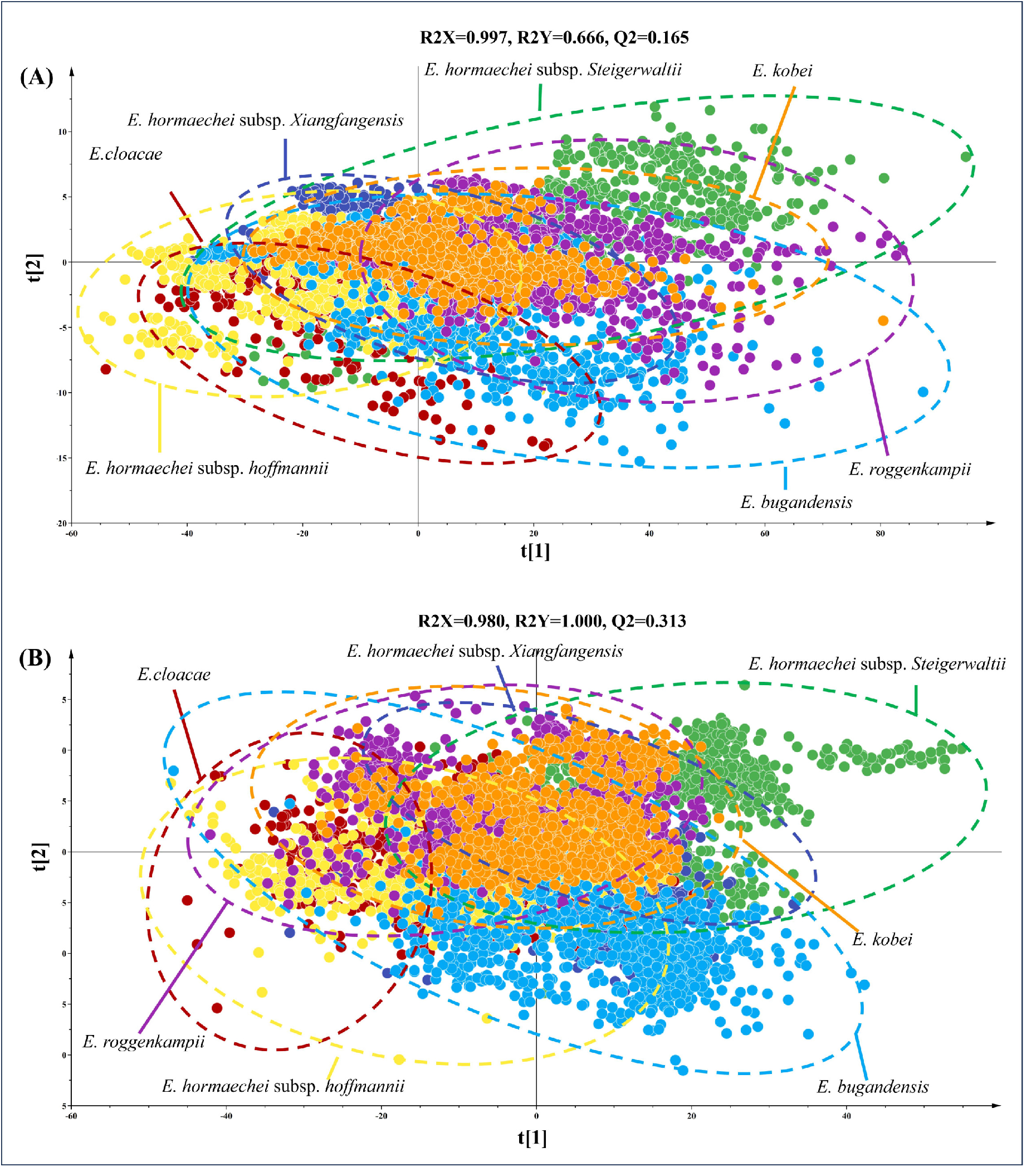
Clustering analysis of SERS spectra of seven ECC species. (A) Scatterplot of SERS spectra via OPLS-DA without normalization. (B) Scatterplot of SERS spectra via OPLS-DA with normalization.

### 3.4 Comparison of supervised learning algorithms

To enable quantitative prediction of ECC, we employed six machine learning algorithms to develop an optimal decision-making model. Before analyzing the SERS data, we present the results of each model’s parameter combinations obtained through grid search using a score gradient plot (Supplementary Table S2). We found that the recognition accuracy of each model was improved with the combination and iteration of parameters. Subsequently, the obtained best parameter combination was inputted into each function, and the models’ performance were evaluated using five different evaluation metrics. As shown in Table 1, the XGBoost algorithm exhibited superior performance with the highest accuracy (accuracy = 97.78%) and stability (5Fold = 97.45%). Notably, Gboost, RF, SVM, and DT also achieved satisfactory results, possibly due to these algorithms belonging to ensemble learning, which possess the ability to handle non-linear relationships and strong feature selection. However, the AdaBoost algorithm failed to differentiate different species of ECC (accuracy = 43.13%), because of the overlapping or intersecting SERS spectra of ECC in feature space, thereby affecting the feature selecting of AdaBoost.

**Table 1.** Comparison of the prediction capabilities of six supervised machine learning models in the analysis of the SERS spectra of *Enterobacter cloacae* complexes.

| Algorithm | Accuracy | Precision | Recall | F1-score | 5Fold |
| --- | --- | --- | --- | --- | --- |
| XGBoost | 97.78% | 97.78% | 97.76% | 97.78% | 97.45% |
| GBoost | 97.49% | 97.49% | 97.44% | 97.49% | 96.67% |
| RF | 91.80% | 91.80% | 91.83% | 91.84% | 92.72% |
| SVM | 90.55% | 90.55% | 90.56% | 90.51% | 89.21% |
| DT | 84.56% | 84.56% | 84.54% | 84.55% | 82.10% |
| AdaBoost | 43.13% | 43.13% | 44.47% | 43.04% | 45.56% |

ROC curve and confusion matrix are commonly used to assess the performance of classification models. We employed the One-vs-All strategy to plot the ROC (Figure 3A) curves for each model, evaluating their ability to discriminate false positive rate (FPR) and true positive rate (TPR) on the test set. The results demonstrate that XGBoost achieved the highest AUC value (AUC=0.9982), while the remaining algorithms also exhibited performance consistent with the metrics provided above. For the optimal classification model, we used the confusion matrix (Figure 3B) to examine the model’s performance on spectral data in detail. It can be seen that XGBoost successfully identified all ECC spectra. For the performance on *E. hormaechei* subsp. *steigerwaltii*, 1.05% of the spectra were incorrectly classified as *E. hormaechei* subsp. *hoffmanniic*. There were also 0.51%, 0.51%, 0.51% of *E. hormaechei* subsp. *xiangfangensis* spectra misidentifying as *E. cloacae, E. hormaechei* subsp. *hoffmanniic* and *E, bugandensis*, respectively. The average recognition accuracy of the XGBoost model was 97.75%, which further demonstrated the potential of this algorithm in distinguishing SERS spectrum of different ECC.

**Figure 3.**
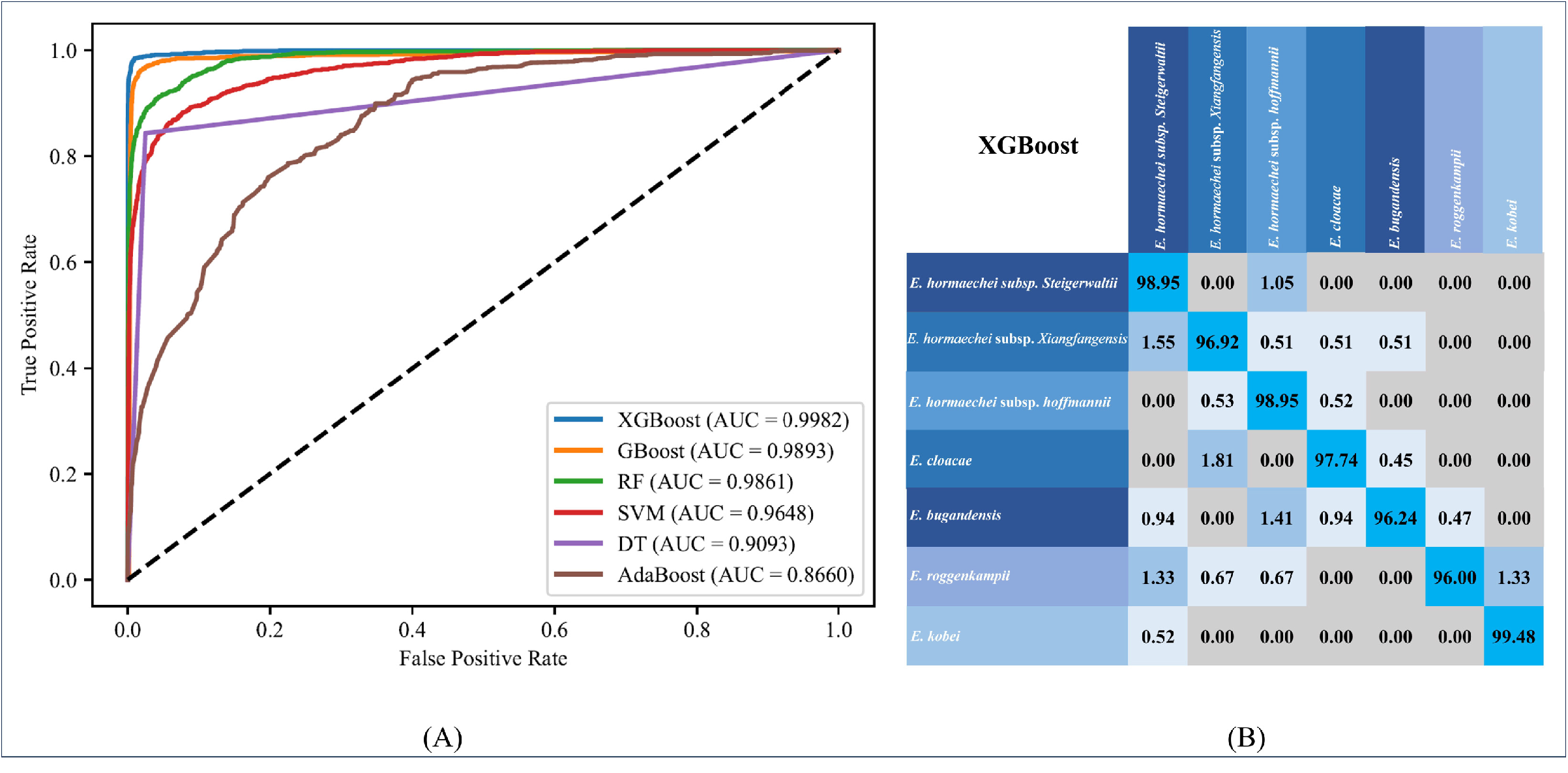
ROC curve and confusion matrix for the machine learning support vector machine model when applied to SERS spectra of ECC. (A) ROC curve. (B) Confusion matrix. The numbers in the confusion matrix represent the possibility of correctly classified (diagonal) or mis-identified (off-diagonal) spectra, respectively.

## Discussion

According to the China Antimicrobial Resistance Surveillance System (CARSS) data, *Enterobacter* spp. rank fifth in the isolation rate of Gram-negative bacteria, accounting for 3-5% of all bacterial isolates in China (Qin et al., 2024). *Enterobacter* spp. isolated from clinical samples are usually reported as *E. cloacae*, and sometimes *E. asburiae, E. hormaechei*, or *E. kobei* by phenotypic methods, all belonging to ECC. Due to the similarities between species, the identification of ECC is often inaccurate and can be variable when repeating the test. However, the discrimination of ECC is clinically important due to the resistance pattern variations among different species. Researchers have shown that most of the carbapenem-resistant isolates were identified as *E. hormaechei subsp*.*xiangfangensis* ST171, a clone circulating globally (Ganbold et al., 2023). Besides, colistin hetero-resistance was found in all or most of *E. roggenkampii, E. kobei, E. chuandaensis* and *E. cloacae* but rarely seen in *E. hormaechei* subspecies and *E. ludwigii* (Fukuzawa et al., 2023). Fortunately, with the development of new technologies, the taxonomy of ECC has evolved over time, from phenotypic methods (Georgiades & Raoult, 2010), such as Gram staining, biochemical assays, and MALDI-TOF MS, to molecular approaches, such as 16S rRNA gene sequencing (Kim et al., 2014), marker gene hsp60 sequencing (Hoffmann & Roggenkamp, 2003) and WGS based ANI and DDH (Goris et al., 2007; Richter & Rossello-Mora, 2009). The species and subspecies within ECC have been assigned to a more detailed classification. To date, twelve species including *E. cloacae, E. hormaechei, E. asburiae, E. cancerogenus, E. kobei, E. ludwigii, E. mori, E. nimipressuralis, E. roggenkampii, E. chengduensis*, and *E. bugandensis* and *E. soli* are assigned to ECC (Fukuzawa et al., 2023; Ganbold et al., 2023). *E. hormaechei* can be subsequently divided into five subspecies (*E. hormaechei* subsp. *steigerwaltii*, subsp. *oharae*, subsp. *xiangfangensis*, subsp. *hoffmannii*, and subsp. *hormaechei*), adding more complexities in the identification of ECC (Qiu et al., 2024).

In clinical laboratories, the accurate identification of ECC species and subspecies still remains a challenge. Routine identification of ECC is mainly dependent on phenotypic characteristics by using commercialized systems, such as Vitek 2 compact and the MALDI-TOF MS technology (De Florio et al., 2018), despite that both methods can only give an ambiguous result. This was also confirmed in our study that Vitek 2 compact failed to assign ECC species and subspecies and Vitek MS only correctly identified one strain of *E. kobei*. Molecular methods are more suitable for precisely identification of the ECC on species level. *Hsp60* typing is the earliest developed and currently most widely used method for this purpose (Hoffmann & Roggenkamp, 2003). However, disadvantages are appearing due to the reclassification of species and subspecies, leading to unclassified or misclassified results by *hsp60*. Recently, multi-plex real-time PCR and combination of single gene (*dnaJ*) real-time PCR plus MALDI-TOF MS for precise ECC identification have been reported (Ji et al., 2021; Pavlovic et al., 2012). These were effective for limited species including *E. cloacae, E. asbuiae, E. hormaechei, E. kobei* and *E. ludwigii* and unable to identify subspecies. So far, only WGS based ANI and DDH are reliable methods for accurate characterization of ECC species and subspecies. Considering the long-period and high-cost of WGS and the difficulty in data analysis, there is a significant requirement to seek easier and cheaper methods.

In recent years, SERS coupled with machine learning algorithms has been emerging as a new technology for the rapid and accurate discrimination of various bacterial pathogens due to its strong Raman effects (Wang et al., 2021). Different SERS substrates and machine learning algorithms have been tried during the exploration of using SERS technique as a quantitative analytical tool for bacterial pathogen diagnosis. Though not developed long, SERS has been successfully applied in the identification of multiple bacteria, like *Escherichia coli, Salmonella typhimurium, Staphylococcus aureus, Staphylococcus epidermidis, Bacillus megaterium* and so on (Rodriguez et al., 2023; Wang et al., 2010; Zhou et al., 2020). Previous studies have demonstrated that simple average SERS spectral analysis plus machine learning algorithms is sufficient for discriminating biological samples based on significant variations in characteristic peaks (Tang, Li, et al., 2022; Wang et al., 2022; Yang et al., 2022). In this study we attempted for the first time to apply SERS in the identification of ECC. Since ECC consists of very much closely related species, it is challenging to accurately distinguish the SERS spectra of different species. To overcome this limitation, deconvoluted SERS spectra were generated to identify subtle molecular vibrations, which has been successfully used to detect differences in very similar species such as Candida and *Shigella* (Liu et al., 2023; Pezzotti et al., 2022). On this basis, we extracted a list of characteristic peaks within ECC. Except for *E. hormaechei subsp. xiangfangensis* and *E. bugandensis* which needs to be identified based on characteristic peaks combination, the other five species all had at least one characteristic peak unique to themselves. However, despite being recognized as unique characteristic peaks, the molecular vibrations were quite close, posing high requirements in detection.

Despite our efforts to minimize undesired effects on Raman spectroscopy measurements during the acquisition of SERS signals, the measured spectral signals still encompass extraneous contributions from the instrument or the sample itself. Hence, data cleaning becomes imperative to eliminate these detrimental effects (Guo et al., 2021). Thus, we processed the spectral feature matrix with maximum and minimum normalization and baseline correction. Subsequently, the processed feature matrices were used for OPLS-DA clustering analyses. This algorithm used specific shapes and combined features to determine sample clusters. The R2X, R2Y, and Q2 metrics were used to appraise ECC species and assess the quality of SERS data. Neverthelss, the high dimensionality and similarity of the SERS spectral data still pose significant challenges for the OPLS-DA clustering algorithm to effectively distinguish different ECC species. Therefore, exploring more advanced methods for spectral data analysis is necessary.

Ciloglu FU et al combined SERS and deep learning techniques for drug-resistant *Staphylococcus aureus* detection, achieving an accuracy of 97.66% and an aAUC of 0.99 (Ciloglu et al., 2021). This breakthrough garnered significant attention and spurred numerous researchers to explore intelligent spectral analysis. In this study, we used normalized and baseline-corrected spectral data as input to construct six ensemble learning models. We employed the GridSearch algorithm to analyze the appropriate hyperparameters of various models (Ma et al., 2023; Tang, Qiao, et al., 2022). The parameter combination yielding the highest final score for each model was selected to examine SERS data and identify different species of ECC in an independent dataset. Among the six models, XGBoost demonstrated the most accurate diagnosis with the highest efficiency in analyzing different species of ECC. This novel diagnostic method holds a promising potential to attain early prediction of ECC, surpassing the capabilities of existing methods.

In summary, SERS coupled with machine learning algorithms showed the potential in the identification of highly similar ECC, enabling us to understand the aspect of clinical significance, epidemiology, and drug resistance of ECC at the species level. In comparison to currently available phenotypic and molecular methods, this method is rapid, accurate, and cost-efficient that is suitable for future use in routine diagnostic tests.

## Acknowledgements

None.

## Fundings

This work was supported by National Science Foundation for Young Scientists of China (82202541), Peking Union Medical College Hospital Talent Cultivation Program Category D (UHB12396) and Fundamental Research Funds for the Central Universities (3332022012).

## Supplementary materials

Supplementary Table S1. Identification results of 92 *Enterobacter cloacae* complexes strains by Vitek 2 Compact, Vitek MS and whole genome sequencing.

Supplementary Table S2. The Best Combination of hyperparameter for Different Machine Learning Algorithms.

